# Genome-wide association study of pain sensitivity assessed by questionnaire and the cold pressor test

**DOI:** 10.1101/837526

**Authors:** Pierre Fontanillas, Achim Kless, 23andMe Research Team, John Bothmer, Joyce Y. Tung

**Affiliations:** 23andMe Inc., Sunnyvale, California, USA; Grünenthal Innovation, Grünenthal GmbH, Aachen, Germany

## Abstract

Altered pain sensitivity is believed to play an important role in the development of chronic pain, a common debilitating condition affecting an estimated 1 in 5 adults. Pain sensitivity varies broadly between individuals, and it is notoriously difficult to measure at scale in large populations. Although pain sensitivity is known to be moderately heritable, only a small number of genetic studies have been published and they have had limited success in identifying the genetic architecture of pain sensitivity. In this study, we deployed an online pain sensitivity questionnaire (PSQ) and an at-home version of the cold pressor test (CPT) in a large genotyped cohort. We performed genome-wide association studies (GWAS) on the PSQ score (n = 25,321) and CPT duration (n = 6,853). Despite a reasonably large sample size, we identified only one genome-wide significant locus associated with the PSQ score, which was located in the *TSSC1* gene (rs58194899, OR = 0.950 [0.933-0.967], P-value = 1.9*10^−8^). This gene is responsible for intracellular cell trafficking and it could modulate pain sensitivity via changes in neuroplasticity. PSQ score is genetically correlated with chronic and acute pain, including chronic neck and shoulder pain (r_g_ = 0.71), fracture (0.71), rheumatoid arthritis (0.68), osteoarthritis (0.38), and pneumothorax (0.82). PSQ score was also genetically correlated with known risk factors, such as the length of working week (0.65), noisy workplace (0.41), smoking (0.36), or extreme BMI (0.23). Gene-based analysis followed by pathway analysis showed that GWAS results were enriched for genes expressed in the brain, predominantly in the frontal cortex and basal ganglia, and enriched for genes involved in neuronal development and glutamatergic synapse signaling pathways. Finally, we confirmed that females with red hair were more sensitive to pain and found that genetic variation in the *MC1R* gene was associated with an increase in pain sensitivity as assessed by the PSQ. Overall, we detailed the genetic background of pain sensitivity using scalable at-home procedures that may serve as a blueprint for larger genetic studies.

**Author Summary:** Despite decades of research and many advances in our understanding of the physical, emotional, and psychological aspects, the genetic contribution to pain sensitivity is still largely unknown. We administered a pain sensitivity questionnaire in parallel with an experimental cold pressor test in a large, genotyped cohort of 31,000 research participants, and found a novel genome-wide genetic association in *TSSC1*. This gene is responsible for intracellular cell trafficking and it could modulate pain sensitivity via changes in neuroplasticity. Overall, the pain sensitivity measurements revealed strong positive genetic correlations with chronic and acute pain conditions, as well as with unhealthy traits and behaviors, such as obesity, smoking, or difficult working conditions. The genetic association results also suggested, perhaps unsurprisingly, that only the brain tissues could be relevant for pain sensitivity, and identified enrichments for glutamatergic synapse pathways. Despite the complexity of pain sensitivity, our study demonstrated that it is now possible to deploy sophisticated pain questionnaires online and perform experimental medicine tests in at-home settings to further study the genetic architecture of pain sensitivity.

## Introduction

It has been established that pain sensitivity is predictive of acute postoperative pain, and of risk for future chronic pain conditions. The assessment of pain sensitivity requires well-controlled experimental pain and emotional stimuli [1]. In general, such assessments are time-consuming in clinical settings because there is substantial inter-individual variability in pain sensitivity and perception [2][3]. As a result, there have been few studies with large sample sizes, impeding progress in understanding the genetic architecture of pain sensitivity. Although hundreds of genes have been proposed to have associations with different types of pain [4], most pain genetics studies analyzed small sample sizes, often using candidate gene or gene panel approaches. To date, the number of genome-wide association studies (GWAS) on pain phenotypes is still very limited. The largest pain GWAS have been performed in the UK Biobank and 23andMe, Inc. cohorts for chronic pain [5][6], knee pain [7], neck and shoulder pain [8], and migraine [9][10]. These studies have identified dozens of putative causal genes, which are primarily expressed within brain tissues and have been implicated in neurogenesis, neuronal development, neural connectivity, and cell-cycle processes. Pain phenotypes have been correlated with a range of psychiatric, personality, autoimmune, anthropometric, and circadian traits. Only a couple of small GWAS studies have directly explored the genetic architecture of pain sensitivity [11][12]. They have found a small number of associations, but none of them have been replicated.

Several clinical and population-based studies have also reported that individuals who naturally have red hair tend to be more resistant to local anesthetics and more sensitive to thermal and dental pain [13]. Red hair, as well as fair skin and freckles, is associated with genetic variations of the melanocortin-1 receptor (MC1R), and it has been suggested that these mutations could directly modulate pain sensitivity, particularly in women [14–16].

We recently validated an online version of the Pain Sensitivity Questionnaire (PSQ) and an at-home version of the cold pressor test (CPT), both of which are used in clinical assessments of pain [17]. The PSQ asks participants to imagine 14 painful situations and 3 non-painful control situations. Subjects are asked to rate their painfulness on a 0–10 numeric scale. The CPT measures how long subjects can immerse one hand in cold ice water. The validation study demonstrated that these two pain sensitivity measures can be consistently collected online and allow pain sensitivity analyses in very large cohorts.

In this study, we performed GWAS on PSQ score and CPT duration in a large European-ancestry cohort (n = 31K) of genotyped individuals, followed by gene-based tests and enrichment analyses. We also attempted to replicate the association between hair color, *MC1R* genetic variants, and pain sensitivity.

## Results

A total of 25,321 and 6,853 research participants of European ancestry were included in the PSQ and CPT GWAS analyses, respectively (Table 1). The two cohorts were largely independent; only 1,534 participants were included in both analyses. The sex ratio was unbalanced in both cohorts, with 71% and 63% of females, respectively. On average, females reported a higher pain sensitivity with a mean PSQ score of 3.23±0.01 versus 3.11±0.01 in males, as well as a lower tolerance to the CPT with 72±0.7 vs. 94±1.0 seconds of hand immersion in ice water. The PSQ and CPT distributions are presented in S1 FIG, and a full description of the overlapping participants has been published elsewhere [18].

**Table 1:** Total number of participants included in the different analyses and mean (and SE) of PSQ score and CPT by sex and age classes.

|  | PSQ (score) |  | CPT (in seconds) |  |
| --- | --- | --- | --- | --- |
|  | Females | Males | Females | Males |
| **Full cohort (used in GWAS):** |  |  |  |  |
| # of participants | *18,060* | *7,261* | *4,343* | *2,510* |
| All | 3.23 (0.01) | 3.11 (0.01) | 72.39 (0.75) | 93.9 (1.01) |
| **Per age classes:** |  |  |  |  |
| <40 years | 3.16 (0.018) | 3.04 (0.028) | 73.38 (1.00) | 94.67 (1.36) |
| 40-60 | 3.27 (0.017) | 3.12 (0.026) | 68.95 (1.42) | 94.72 (1.81) |
| >60 | 3.25 (0.017) | 3.16 (0.024) | 74.83 (1.91) | 89.46 (2.71) |
| **MC1R carriers:** |  |  |  |  |
| Non-carrier | 3.22 (0.012) | 3.10 (0.018) | 71.98 (0.91) | 92.70 (1.19) |
| Carrier1 | 3.26 (0.019) | 3.14 (0.029) | 73.51 (1.42) | 96.98 (2.01) |
| Carrier2+ | 3.33 (0.062) | 3.18 (0.093) | 72.36 (4.39) | 100.35 (6.18) |
| **Hair color cohort:** |  |  |  |  |
| # of participants with hair color information | *10,928* | *4,239* | *3,375* | *1,868* |
| Proportion of redhead | *5.1%* | *3.2%* | *5%* | *3.8%* |
| Not redhead | 3.22 (0.013) | 3.13 (0.02) | 72.12 (0.88) | 92.22 (1.20) |
| Redhead | 3.37 (0.060) | 3.01 (0.10) | 71.69 (3.77) | 100.43 (5.77) |

The GWAS on PSQ score produced one locus that reached genome-wide significance. The lead SNP (rs58194899: OR=0.950 [95%CI 0.933-0.967], P-value = 1.9×10^−8^, MAF = 0.47; Table 2) was located in the *TSSC1* gene, which is also known as *EIPR1* (Fig 1A and Fig 2B). The associated haplotype was relatively small (54 variants in the 99% credible set), and entirely located within the *TSSC1* gene boundary (Figure 2A). To date, no studies have reported a phenotypic association with this haplotype (https://genetics.opentargets.org/). However, a cognitive decline GWAS reported an independent association in the *TSSC1* gene (lead variant rs75365287) [19]. TSSC1 is a component of the endosomal retrieval pathway. It plays a critical role as a regulator of both Golgi-associated retrograde protein (GARP) and endosome-associated recycling protein (EARP) functions, as well as the transport of internalized proteins to the plasma membrane [20,21]. TSSC1 and the GARP/EARP complexes are not known to be involved in pain traits. However, TSSC1 is overexpressed in the brain, particularly in the hypothalamus, and in the frontal and interior cingulate cortices (GTEx v8).

**Table 2:** Top GWAS variants and MCR1 association results for PSQ and CPT traits. PSQ score were analyzed using a linear model. CPT duration (in seconds) were analyzed using a survival model.

| *CHRPOS* |  | *ID* | *Alleles* | *MAF* | *Gene context* | PSQ GWAS |  |  | CPT GWAS |  |  |
| --- | --- | --- | --- | --- | --- | --- | --- | --- | --- | --- | --- |
|  |  |  |  |  |  | *Effect* | *SE* | *P-value* | *Effect* | *SE* | *P-value* |
| Top PSQ GWAS Variants: |  |  |  |  |  |  |  |  |  |  |  |
| 2 | 3280983 | rs58194899 | G/A | 0.47 | TSSC1 | -0.051 | 0.009 | 1.9e-08 | -0.01 | 0.018 | 5.8e-01 |
| 18 | 10008669 | rs142738119 | T/C | 0.001 | VAPA, APCDD1 | 0.696 | 0.138 | 4.7e-07 | -0.035 | 0.275 | 9.0e-01 |
| 13 | 101893919 | rs12583902 | G/A | 0.166 | NALCN | 0.059 | 0.012 | 7.6e-07 | -0.025 | 0.023 | 2.9e-01 |
| Top CPT GWAS Variant: |  |  |  |  |  |  |  |  |  |  |  |
| 17 | 65520139 | rs141828201 | G/A | 0.027 | PITPNC1 | 0.036 | 0.031 | 2.6e-01 | 0.356 | 0.066 | 2.4e-07 |
| MC1R Variants: |  |  |  |  |  |  |  |  |  |  |  |
| 16 | 89986144 | rs1805008 | C/T | 0.071 | MC1R | 0.031 | 0.017 | 6.8e-02 | -0.032 | 0.034 | 3.5e-01 |
| 16 | 89986117 | rs1805007 | C/T | 0.075 | MC1R | 0.047 | 0.017 | 5.1e-03 | -0.039 | 0.033 | 2.4e-01 |
| 16 | 89986546 | rs1805009 | G/C | 0.019 | MC1R | 0.041 | 0.032 | 2.0e-01 | -0.089 | 0.063 | 1.5e-01 |

**Figure 1:**
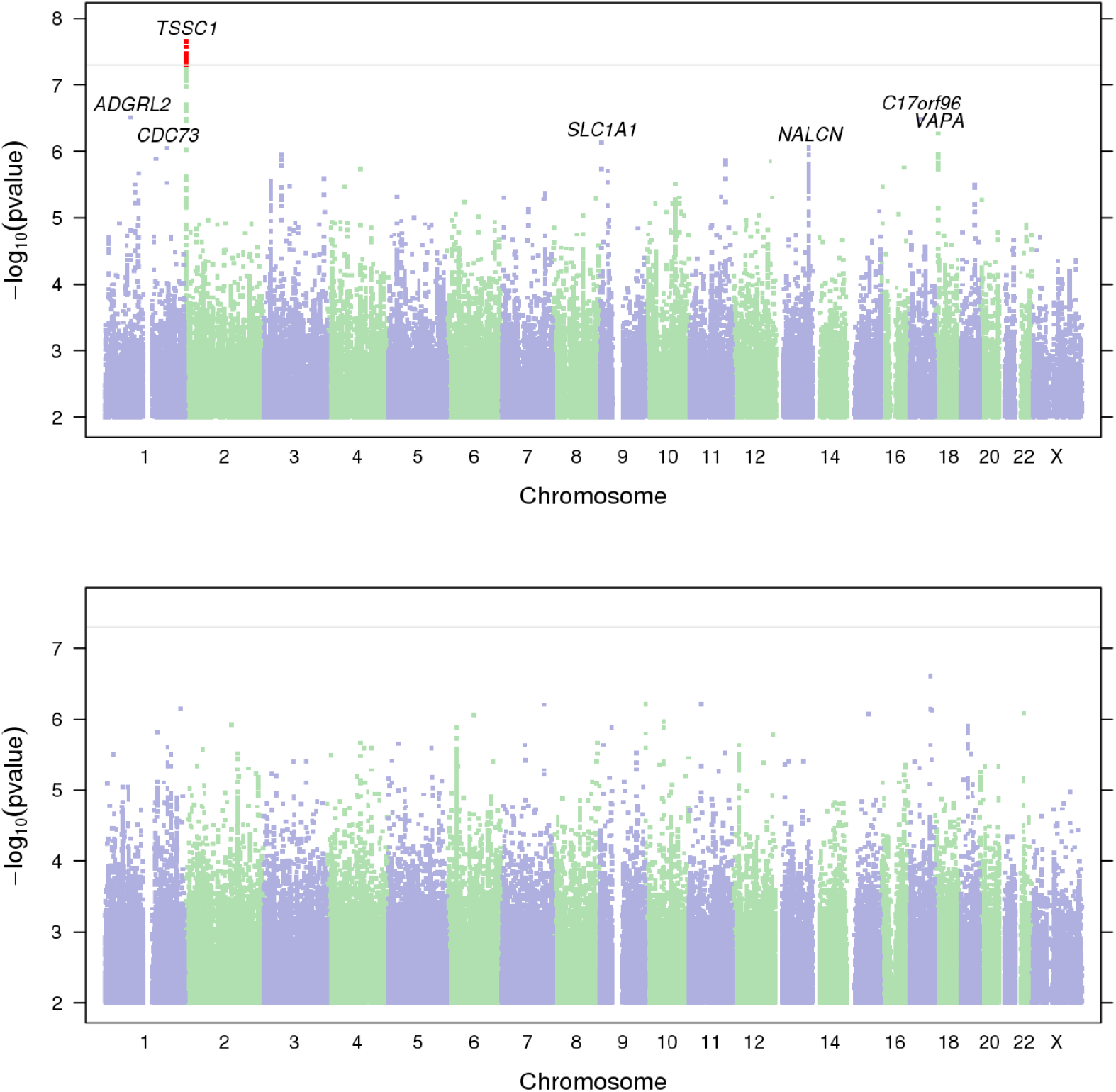
Manhattan plots for PSQ and CPT GWAS.

**Figure 2:**
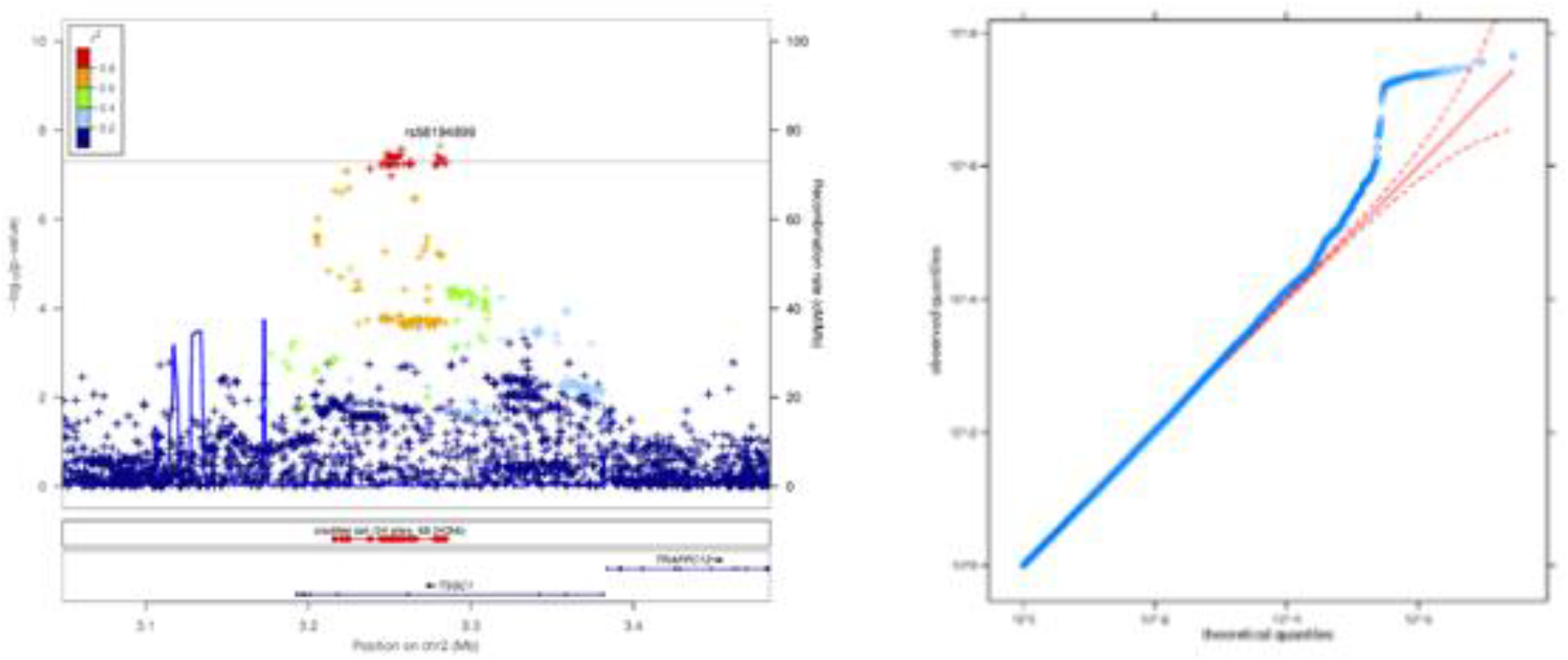
Regional association plot for TSSC1 locus and QQ plot from the PSQ GWAS

Using the PSQ GWAS summary statistics, we performed a gene-based association analysis in MAGMA (v1.07), followed by a gene-set enrichment analysis in FUMA (GENE2FUNC, v1.3.5). A total of 58 genes were identified with an adjusted P-value < 1.0×10^−4^ (Table 3). These genes were over-expressed in the brain, especially in the frontal cortex, basal ganglia, amygdala, and hypothalamus (S5 Fig). They were significantly enriched for genes involved in brain development and synaptic signaling pathways (Table 3 and S6 Table). Like *TSSC1*, many of these genes are involved in Golgi apparatus function (*RBFOX1, PARK2, WWOX, PRKG1, LARGE, GPC6, PCSK6, HS3ST4, WWOX, SLC39A11, PRKCE, TENM2*, and *CNTNAP2*; S6 Fig). Several genes identified by MAGMA are active in glutamatergic synapses, which are involved in pain sensation and transmission (*PTPRD, NRG1, NRG3, DLG2, GPC6*, and *GRID2*). Finally, among the genes that were most strongly associated with PSQ score, *CSMD1*, *LRP1B*, and *DMD* were not known to be directly involved in pain sensitivity but were linked to neurological diseases like bipolar disorder. The traits with the highest genetic correlations with PSQ, as computed on LD Hub, are listed in Table 4. PSQ score was strongly genetically correlated with chronic pain related phenotypes, such as neck-and-shoulder pain (r_g_ = 0.71), rheumatoid arthritis (0.68), or mononeuropathies (0.53), and with acute pain phenotypes, such as pneumothorax (0.82) or fracture (0.71). It was also strongly correlated with health risk factors and behaviors, such as the length of working week (0.65), working in a noisy environment (0.42), shift work (0.41), smoking (0.36), and extreme BMI (0.23). We also observed strong positive genetic correlation between the PSQ score and ADHD (0.67), but negative correlations with schizophrenia (−0.21), bipolar disorder (−0.25), and neuroticism (−0.22).

**Table 3:**
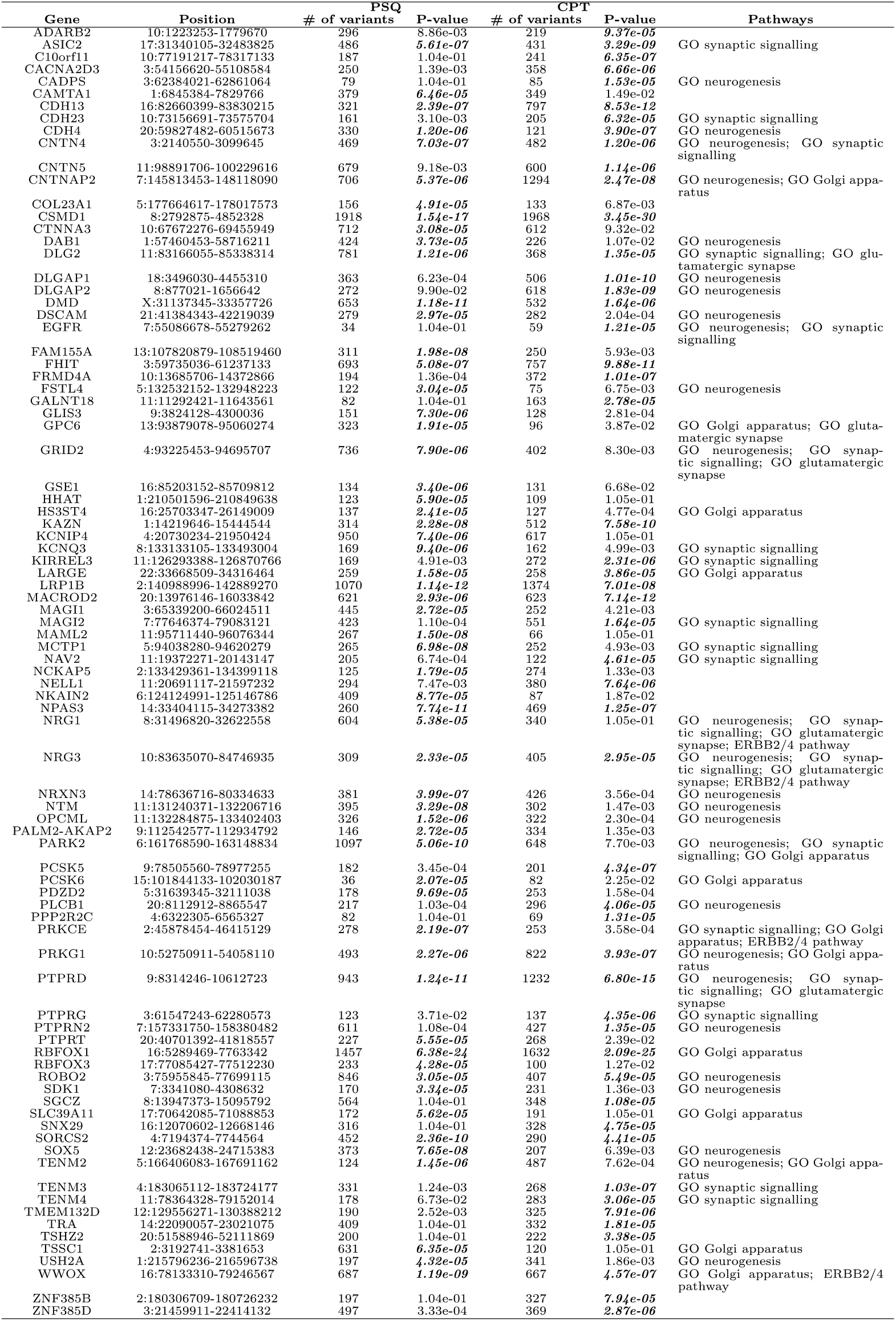
Gene-based (MAGMA) and pathway analysis results for PSQ and CPT. A total of 58 and 50 significantly associated genes (P-value < 10^−4^, highlighted in bold) for PSQ and CPT, respectively, including 21 genes identified in both pain sensitivity measures.

**Table 4:** Top genetic correlations between PSQ and LD Hub phenotypes

| Trait | rG | rG se | z | P-value | h2 | h2 se |
| --- | --- | --- | --- | --- | --- | --- |
| Pneumothorax | 0.8217 | 0.4166 | -1.9722 | 0.0486 | 0.0031 | 0.0014 |
| Fracture resulting from simple fall | 0.7123 | 0.3442 | -2.0692 | 0.0385 | 0.0518 | 0.0154 |
| Neck/shoulder pain for 3+ months | 0.7118 | 0.3416 | 2.0835 | 0.0372 | 0.0214 | 0.0064 |
| Rheumatoid arthritis | 0.6804 | 0.3244 | 2.0974 | 0.036 | 0.0044 | 0.0014 |
| ADHD | 0.6738 | 0.3204 | 2.1031 | 0.0355 | 0.2428 | 0.0988 |
| Length of working week for main job | 0.6473 | 0.2681 | 2.4141 | 0.0158 | 0.0191 | 0.0029 |
| Nasal polyps | 0.6148 | 0.3046 | -2.0187 | 0.0435 | 0.0053 | 0.0016 |
| Complications of internal orthopaedic prosthetic devices | 0.614 | 0.3208 | 1.9138 | 0.0557 | 0.0052 | 0.0017 |
| Cholelithiasis/gall stones | 0.5583 | 0.2568 | 2.1743 | 0.0297 | 0.0083 | 0.0015 |
| Mononeuropathies of upper limb | 0.5271 | 0.2086 | 2.5264 | 0.0115 | 0.0131 | 0.0018 |
| Internal derangement of knee | 0.4772 | 0.231 | 2.0662 | 0.0388 | 0.0081 | 0.002 |
| Duration of walks | 0.4281 | 0.1406 | 3.0447 | 0.0023 | 0.0463 | 0.0029 |
| Heavy physical activity (eg: weeding, carpentry) | 0.4168 | 0.1579 | 2.6392 | 0.0083 | 0.0343 | 0.002 |
| Noisy workplace | 0.4153 | 0.1782 | 2.3303 | 0.0198 | 0.062 | 0.006 |
| Job involves shift work | 0.4098 | 0.1847 | 2.2185 | 0.0265 | 0.03 | 0.0029 |
| Osteoarthritis | 0.3844 | 0.1659 | 2.3169 | 0.0205 | 0.0186 | 0.0018 |
| Disability diagnosed by doctor | 0.3595 | 0.1612 | 2.23 | 0.0257 | 0.025 | 0.0019 |
| Number of cigarettes previously smoked daily | 0.3557 | 0.165 | 2.1554 | 0.0311 | 0.0996 | 0.0143 |
| Falls in the last year | 0.3544 | 0.1513 | 2.3416 | 0.0192 | 0.0321 | 0.002 |
| Getting up in morning | 0.3128 | 0.1151 | 2.7181 | 0.0066 | 0.0686 | 0.0031 |
| Mouth/teeth dental problems (Dentures) | 0.3069 | 0.1363 | 2.2521 | 0.0243 | 0.048 | 0.0028 |
| Pack years of smoking | 0.2538 | 0.1376 | 1.8445 | 0.0651 | 0.1086 | 0.0104 |
| Risk taking | 0.247 | 0.1176 | 2.1013 | 0.0356 | 0.0561 | 0.0028 |
| Extreme bmi | 0.2348 | 0.1271 | 1.8468 | 0.0648 | 0.6852 | 0.0576 |
| Overweight | 0.221 | 0.1114 | 1.9843 | 0.0472 | 0.1093 | 0.0068 |
| Morning/evening person | -0.1871 | 0.1022 | -1.83 | 0.0672 | 0.1157 | 0.0046 |
| Schizophrenia | -0.2109 | 0.0886 | -2.3805 | 0.0173 | 0.462 | 0.0192 |
| Neuroticism score | -0.2165 | 0.1132 | -1.9126 | 0.0558 | 0.1191 | 0.0064 |
| Guilty feelings | -0.2183 | 0.122 | -1.7895 | 0.0735 | 0.052 | 0.0028 |
| Coronary artery disease | -0.2341 | 0.1325 | -1.7672 | 0.0772 | 0.0792 | 0.0058 |
| Bipolar disorder | -0.2473 | 0.1442 | -1.7147 | 0.0864 | 0.4282 | 0.0362 |
| Suffer from nerves | -0.2499 | 0.1295 | -1.9297 | 0.0536 | 0.046 | 0.0031 |
| Narcolepsy | -0.2509 | 0.1315 | -1.9081 | 0.0564 | 0.049 | 0.0027 |
| Time spent watching television | -0.2701 | 0.1257 | -2.148 | 0.0317 | 0.0991 | 0.004 |
| Nervous feelings | -0.2843 | 0.1283 | -2.2158 | 0.0267 | 0.0669 | 0.0042 |
| Psoriasis | -0.4522 | 0.2555 | -1.7702 | 0.0767 | 0.0075 | 0.0017 |
| Celiac disease | -0.5604 | 0.2962 | -1.8918 | 0.0585 | 0.2959 | 0.0502 |
| Acetate | -0.5861 | 0.3195 | -1.8344 | 0.0666 | 0.0556 | 0.0192 |

The GWAS on CPT duration was underpowered and did not produce any significant genome-wide associations (Fig. 1b, S2 Fig., and S1 Table). Nevertheless, the genetic correlation between PSQ score and CPT duration was *r_g_* = −0.64 ± 0.37 (P-value = 0.088), and the results from the CPT gene-based analysis and pathway analysis were consistent with the PSQ results. Among the 50 CPT genes identified by MAGMA, 21 were also identified in the PSQ analysis (Table 3). Similarly, these 50 genes were overexpressed in brain (S7 Fig.) and enriched for brain development and synaptic signaling pathways (S6 Table).

We also specifically focused on the association results for genes involved in nociception. We obtained a list of 26 nociception genes from the Human Pain Genes Database ([22], https://humanpaingenetics.org/hpgdb/). None of the 21 nociception genes showed evidence of association with PSQ and CPT (S7 Table, MAGMA analysis).

Finally, we assessed the relationship between both pain sensitivity measures, PSQ and CPT, and hair color on a subset of the PSQ and CPT cohorts with available hair color data (Table 1). A total of 15,167 and 5,243 participants were included in the analyses for PSQ score and CPT duration, respectively. Using a Gaussian linear model including age, sex, and the first five genetic principal components as covariables, we showed that participants with red hair reported significantly higher PSQ scores than participants with light blond, dark blond, light brown, dark brown, or black hair (Fig. 3c and S2 Table). Furthermore, females with red hair reported on average higher PSQ scores than non-red hair females, and red or non-red hair males (sex-by-red hair interaction P-value = 0.046; S2 Table and Fig. 3b). We did not observe significant PSQ differences between red and non-red hair males. For CPT, we did not observe any significant associations using a Cox proportional hazards model (S3 Table and S4 FIG.) or an ANOVA on CPT duration converted in ranks (S4 Table). Since red hair color is partially determined by recessive genetic polymorphism in the *MC1R* gene, we explored the association of the three main variants rs1805009 (D294H), rs1805008 (R160W) and rs1805007 (R151C) in the PSQ and CPT GWAS (Table 2 and S1 Table). None of these individual variants passed the genome-wide significant threshold in the GWAS analyses (P-values > 5.1 × 10^−3^). We combined these three variants and defined three categories of *MC1R* variant carriers, Non-carrier (0 *MC1R* recessive allele), Carrier1 (1 allele), and Carrier2+ (>1 alleles), and tested their association with PSQ score and CPT duration. *MC1R* carriers reported significantly higher PSQ score that non-carriers (P-value = 6.8 × 10^−3^ and 1.5 × 10^−2^, respectively; S2 table and Fig. 3a). On the other hand, we did not observe any significant associations between *MC1R* carriers and CPT duration (S3 and S4 Tables, and S3 Fig.)

**Figure 3:**
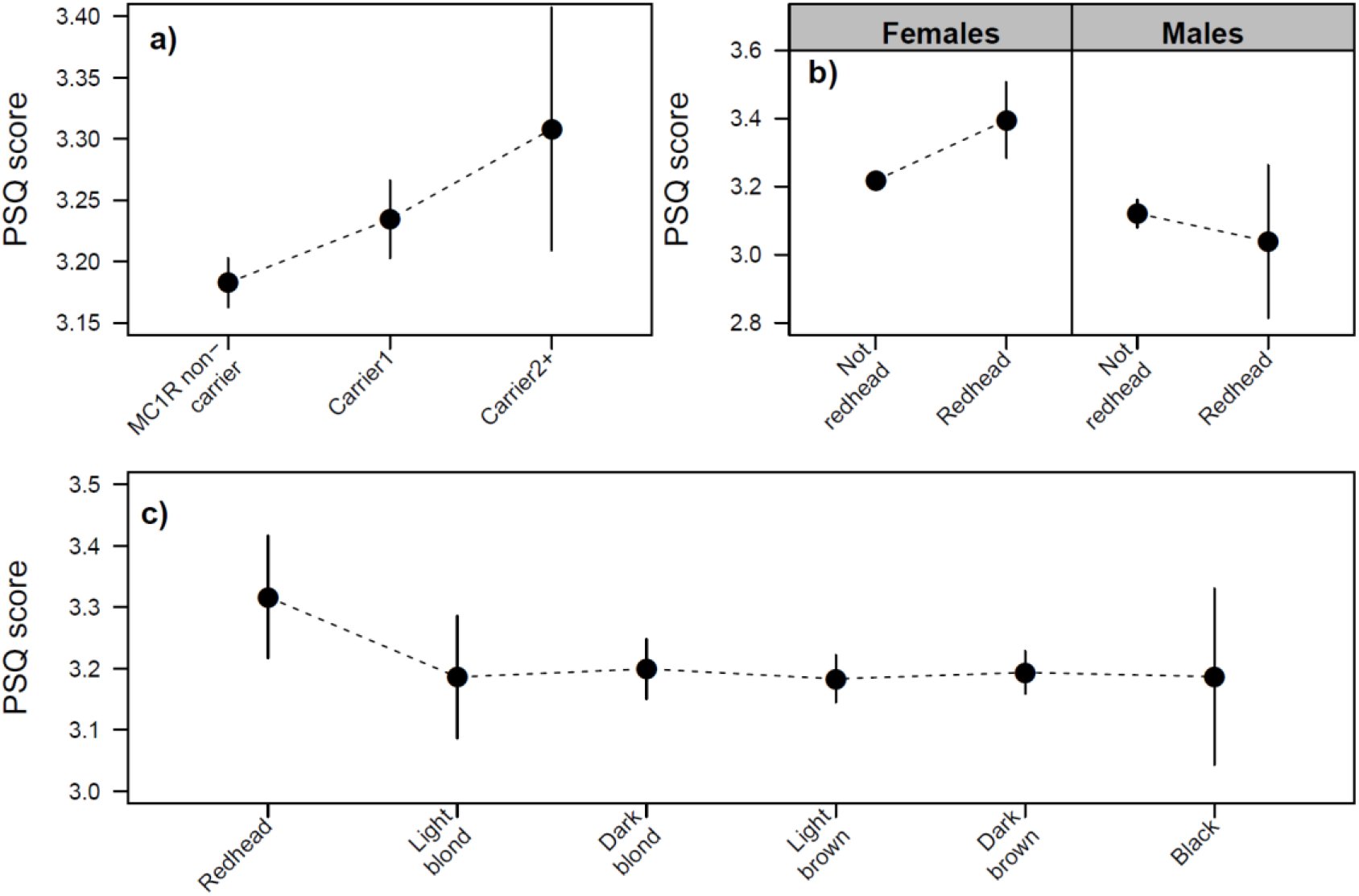
Results of MC1R and hair color association with PSQ score.

## Discussion

The purpose of this study was to identify genetic factors contributing to the individual perception of pain [23]. We used two self-administered pain sensitivity measurements, the PSQ and CPT, and established that they could be reliable phenotypes for evaluating the genetic architecture of pain sensitivity. PSQ score and CPT duration have been previously shown to be only moderately phenotypically correlated (r = −0.22 [−0.27, −0.17]) [8]. The PSQ is a pain intensity rating of imagined painful situations occurring in daily life, whereas the CPT directly measures pain tolerance. Although the PSQ was not designed to cover all dimensions of pain experience, it incorporates the emotional and cognitive components of the pain sensitivity [24].

Most of the genes identified by the association analyses are overexpressed in the brain. This is especially true for the amygdala, the emotional pain processing center, but other components of the pain matrix (e.g. frontal cortex, basal ganglia, and hypothalamus) also showed significant enrichment [25,26]. We found that no brain area seemed to be selectively and exclusively associated with pain sensitivity. These genetic results were in line with recent brain imaging studies that suggested that the individual variability in pain sensitivity is most probably produced by the connectivity of multiple brain areas [27,28]. It is interesting to notice the absence of association enrichments in non-brain tissues. According to the current multifaceted experience concept, pain results from the integration of nociception and the cognitive-emotional state of the individual. Nociceptors are found in skin and mucosa, as well as in a variety of organs, such as the digestive tract, the bladder, the gut, and muscles. However, our analysis of the 21 nociceptor genes identified in HPGDB showed no evidence that genetic polymorphism near these genes were associated with PSQ score or CPT duration. The CPT was designed to quantify the evoked pain and signal sensory responses of cold pain sensitivity. Although underpowered, analyses showed very similar enrichment patterns than the PSQ. Among the combined 87 genes identified by the PSQ and CPT gene-based analyses, ten of them were present in HPGDB (*ADARB2, CACNA2D3, CTNNA3, DMD, KCNQ3, PARK2, PCSK6, PRKG1, PTPRD*, and *TRA*). The vast majority were identified in various migraine GWAS [9]. Pathway enrichment analyses showed that many of these 87 genes are involved in neuron and brain development, and neuron signaling. In particular, they highlighted genes active in glutamatergic synapses. Glutamate receptors have a leading role in pain signal transmission and are often considered promising targets for the treatment of chronic pain [29]. Among the six genes identified in this pathway by PSQ associations, *NGR1* and *NGR3* were already known to be linked to pain sensitivity. Both are involved in the ErbB2 and ErbB4 signaling pathways, which have repeatedly been demonstrated to directly contribute to the development of neuropathic pain [30]. To our knowledge, however, our study is the first where genetic polymorphism in these two genes has been associated with pain sensitivity measurements. On the other hand, the four other genes in the pathway, *PTPRD, DLG2, GPC6*, and *GRID2*, have been all associated with diverse neurological disorders [31–34]. This observation supports earlier findings that neurological disorders and pain sensitivity are intimately linked [35]. The genome-wide significant locus, *TSSC1,* has not previously been associated with pain traits. However, it has been associated with cognitive decline [19], and was recently identified in a study that examined alterations in the postsynaptic protein profile as a consequence of prolonged exposure to morphine [36].

Finally, we confirmed that women with red hair are more sensitive to pain, but we did not observe this relationship in men. None of the three main variants in *MC1R* that control red hair color showed genome-wide significant associations with PSQ score or CPT duration. Nevertheless, in combination, individuals carrying one or more copies of these three variants reported a significant higher pain sensitivity.

Overall, the PSQ score and CPT duration had similar genetic correlation profiles — both were correlated with general pain phenotypes such as neck-and-shoulder pain, rheumatoid arthritis, osteoarthritis, neuropathy, and fracture. We also observed positive genetic correlations between pain sensitivity and disability phenotypes, including complications associated with prosthetic devices. Given that epidemiological studies have regularly reported that unhealthy lifestyle practices accentuate pain sensitivity and chronic pain [37], it was encouraging to observe that the pain sensitivity phenotypes were genetically correlated with unhealthy behaviors such as smoking, extreme BMI, length of working week, shift work, or working in a noisy environment. As expected, we also observed significant genetic correlations between pain sensitivity and neurological disorders or personality traits. Notably, pain sensitivity showed a strong positive genetic correlation with ADHD. This was somewhat unexpected based on the observed genetic architectures described in recent pain and ADHD GWAS studies [38]. On the other hand, we observed negative genetic correlations between pain sensitivity and schizophrenia, neuroticism, and bipolar disorder. It is also interesting to note the absence of genetic correlation with migraine and depression despite the fact that migraine, in particular, is associated with intense pain.

Because the inter-individual variation of pain sensitivity depends on the integration of the sensory pathways and the emotional-cognitive states of individuals, it will require more advanced measures of the full pain sensitivity spectrum to identify and disentangle the genetic architecture of pain sensitivity [39, 40]. This study focused only on a narrow set of pain sensitivity measurements. However, we demonstrated that it is now possible to deploy and collect relevant data at a population scale for fairly sophisticated tests such as the CPT. Large sample sizes are commonly needed for uncovering the genetic architecture of complex traits and our results confirmed that pain sensitivity will not be an exception. A larger PSQ dataset should be particularly informative, not only for validating some of the results of this study, but also for evaluating the potential genetic basis of pain sensitivity differences between females and males, and for focusing on specific imaginary pain situations included in the PSQ.

## Materials and Methods

### Study sample

All participants included in the analyses were drawn from the research participant base of 23andMe, Inc., a personal genetics company. Participants provided informed consent and participated in the research online, under a protocol approved by the external AAHRPP-accredited IRB, Ethical & Independent Review Services (E&I Review). (http://www.eandireview.com; OHRP/FDA registration number IRB00007807, study number 10044-11). We restricted all analyses to a set of unrelated participants having >97% European ancestry, as determined through an analysis of local ancestry. Participants were labelled as related if they shared more than 700 cM of identity-by-descent.

### Pain sensitivity traits

For the assessment of the pain interpretation and perception, we used a pain sensitivity questionnaire (PSQ) and an at-home version of cold pressor test (CPT) on two subsets of 23andMe research participants who self-reported chronic pain conditions [2]. The PSQ is an English-language version of the Pain Sensitivity Questionnaire [3] [41], supplemented with additional questions about the participant’s own memory of painful experiences. The PSQ contains 14 questions in which you should imagine yourself in certain situations. You should then grade how painful they would be, from 0 that stands for no pain to 10, the most severe pain that you can imagine or consider possible. The total PSQ score is the mean of the 14 responses. We also computed two PSQ subscales: PSQ-minor score based on the least painful questions (#14, 3, 6, 12, 11, 10, and 7, ordered from least to most painful), and PSQ-moderate score (#8, 15, 2, 16, 17, 1, 4). For the CPT, participants were asked to prepare their own bath of ice water at home, and to keep their non-dominant hand submerged to the wrist for no more than 150 seconds. A separate consent for the CPT was used: participants reporting neurological or temperature-triggered conditions (e.g. migraine, history of syncope, or Raynaud’s phenomenon) or current injuries to their non-dominant hands at the time of recruitment were ineligible. For additional information on the PSQ and CPT, in particular the validity of these approaches for estimating pain sensitivity [18].

### Genotyping and variant imputation

DNA extraction and genotyping were performed on saliva samples by LabCorp, Inc. Participants were genotyped on one of five Illumina genotyping platforms, containing between 550,000 to 950,000 variants, for a total of 1.6 million of genotyped variants. Samples that failed to reach 98.5% call rate were re-analyzed. Genotyping quality controls included discarding variants with a Hardy-Weinberg *P*<10^−20^, a call rate of <90%, or batch effects. About 57.5M of variants were then imputed against a single unified imputation reference panel, combining the May 2015 release of the 1000 Genomes Phase 3 haplotypes with the UK10K imputation reference panel. Principal components were computed using ~65,000 high quality genotyped variants present in all five genotyping platforms. For more details on genotyping, imputation process, and variant quality controls, see [21].

### GWAS analysis

Imputed dosages and genotyped data were both tested for association with PSQ score or CPT duration (between 0 and 150 seconds). The PSQ scores were inverse normalized and analyzed using a gaussian linear model. The association P-value were computed using a likelihood ratio test. The CPT duration was analyzed using a cox proportional hazards model, a survival model on the CPT time. We included covariates for age, gender, genotyping platform, and the top five principal components to account for residual population stratification. The PSQ association model did not include platform covariables because PSQ participants were all genotyped on platform v4. Results for the X chromosome were computed similarly, with males coded as if they were homozygous diploid for the observed allele. A total of 1.3M genotyped and 25.5M imputed variants passed the pre- and post GWAS quality controls. We furthermore filtered out variants with MAF < 0.1%, which are extremely sensitive to quantitative trait over-dispersion, reducing to 13.7M variants available for follow-up analyses. A detailed description of the variant quality control and GWAS methods can be found in [42]. The full GWAS summary statistics for the 23andMe discovery data set will be made available through 23andMe to qualified researchers under an agreement with 23andMe that protects the privacy of the 23andMe participants. Please visit https://research.23andme.com/dataset-access/ for more information and to apply to access the data.

### MC1R and hair color

We defined three categories of *MC1R* variant carriers by combining three variants, rs1805007, rs1805008, and rs1805009: Non-carrier (0 *MC1R* alleles), Carrier1 (1 allele), and Carrier2+ (>1 alleles) [17]. A self-reported hair color phenotype was available for 63% of the participants in the cohort [43]. It contained six hair color categories: red, light blond, dark blond, light brown, dark brown, and black. We also built a binary red hair variable from this categorical hair color phenotype. Association of MC1R carrier, hair color, and red variables were tested against PSQ score or CPT duration, using gaussian linear and cox proportional hazards models, respectively. We also converted CPT duration in ranks, and tested associations using ANOVAs. The same set of covariables used in GWAS was included in these models.

### Genetic correlation, gene-based and pathway analyses

Genetic correlations between PSQ and a broad list of diseases and traits, were estimated with LD Hub v1.9.1 (http://ldsc.broadinstitute.org/ldhub/), using the default analysis parameters. The gene-based analysis was performed on MAGMA (v1.07) [44]. After correction with a Benjamini-Hochberg procedure, genes with adjusted P-values < 10^−4^ were selected and used in pathway analyses performed on FUMA (GENE2FUNC, v1.3.5; https://fuma.ctglab.nl/). Pathway enrichment was tested for different gene sets, including canonical pathways, Reactome, and GO biological processes. Tissue specificity for the set of selected genes was tested using an enrichment analysis of differentially expressed genes (DEG).

## Acknowledgments

This research was conducted by 23andMe and Grünenthal GmbH. All authors are employees of these companies. The funders provided support in the form of salaries for all authors but did not have any additional role in the study design, data collection and analysis, decision to publish, or preparation of the manuscript. The authors like to thank the contributions from the 23andMe Research Team (Michelle Agee, Adam Auton, Robert K. Bell, Katarzyna Bryc, Sarah L. Elson, Pierre Fontanillas, Nicholas A. Furlotte, David A. Hinds, Naomi Iwata, Jennifer C. McCreight, Karen E. Huber, Aaron Kleinman, Nadia K. Litterman, Joanna L. Mountain, Elizabeth S. Noblin, Carrie A.M. Northover, Steven J. Pitts, J. Fah Sathirapongsasuti, Olga V. Sazonova, Anjali J. Shastri, Janie F. Shelton, Suyash Shringarpure, Chao Tian, Vladimir Vacic, Catherine Weldon, Keng-Han Lin, Yunxuan Jiang, Kimberly McManus, David Poznik, Ethan Jewett, Xin Wang, Barry Hicks).

We thank 23andMe research participants and employees for making this work possible.

Pierre Fontanillas and Joyce Tung are employed by and hold stock or stock options in 23andMe, Inc

## Supporting information

**S1 table:** Top GWAS and MCR1 association results for PSQ and CPT traits. PSQ score were analyzed using a linear model. CPT duration (in seconds) were converted into three categories (δ50s; δ100s; and δ150s), and analyzed using a linear model.

|  |  |  |  |  |  | PSQ GWAS |  |  | CPT (CAT.) GWAS |  |  |
| --- | --- | --- | --- | --- | --- | --- | --- | --- | --- | --- | --- |
| CHR | POS | ID | Alleles | MAF | Gene context | Effect | SE | P-value | Effect | SE | P-value |
| Top PSQ GWAS Variants: |  |  |  |  |  |  |  |  |  |  |  |
| 2 | 3280983 | rs58194899 | G/A | 0.47 | TSSC1 | -0.051 | 0.009 | 1.9e-08 | 0.005 | 0.015 | 7.2e-01 |
| 18 | 10008669 | rs142738119 | T/C | 0.001 | VAPA,APCDD1 | 0.696 | 0.138 | 4.7e-07 | 0.343 | 0.235 | 1.4e-01 |
| 13 | 101893919 | rs12583902 | G/A | 0.166 | NALCN | 0.059 | 0.012 | 7.6e-07 | 0.012 | 0.02 | 5.3e-01 |
| Top CPT GWAS Variants: |  |  |  |  |  |  |  |  |  |  |  |
| 17 | 65520139 | rs141828201 | G/A | 0.027 | PITPNC1 | 0.036 | 0.031 | 2.6e-01 | -0.203 | 0.056 | 2.7e-04 |
| MC1R Variants: |  |  |  |  |  |  |  |  |  |  |  |
| 16 | 89986144 | rs1805008 | C/T | 0.071 | MC1R | 0.031 | 0.017 | 6.8e-02 | 0.011 | 0.029 | 7.0e-01 |
| 16 | 89986117 | rs1805007 | C/T | 0.075 | MC1R | 0.047 | 0.017 | 5.1e-03 | 0.04 | 0.028 | 1.5e-01 |
| 16 | 89986546 | rs1805009 | G/C | 0.019 | MC1R | 0.041 | 0.032 | 2.0e-01 | 0.061 | 0.052 | 2.4e-01 |

**S2 table:** Linear models on PSQ score analyzing MC1R carriers and hair colors.

|  | MC1R model |  |  |  | Hair color model |  |  |  | Redhead x Sex model |  |  |  |
| --- | --- | --- | --- | --- | --- | --- | --- | --- | --- | --- | --- | --- |
|  | Beta | SE beta | t value | P-value | Beta | SE beta | t value | P-value | Beta | SE beta | t value | P-value |
| Intercept | 2.9708 | 0.0330 | 89.94 | **0.00e+00** | 2.9736 | 0.0415 | 71.72 | **0.00e+00** | 3.0909 | 0.0656 | 47.14 | **0.00e+00** |
| Age | 0.0025 | 0.0005 | 4.71 | **2.51e-06** | 0.0031 | 0.0007 | 4.59 | **4.56e-06** | 0.0031 | 0.0007 | 4.62 | **3.80e-06** |
| Sex (Female) | 0.1283 | 0.0188 | 6.84 | **8.37e-12** | 0.0966 | 0.0246 | 3.92 | **8.77e-05** | 0.1051 | 0.0244 | 4.30 | **1.68e-05** |
| PC1 | -2.2823 | 0.2112 | -10.81 | **3.70e-27** | -2.3501 | 0.2709 | -8.68 | **4.52e-18** | -2.3482 | 0.2747 | -8.55 | **1.37e-17** |
| PC2 | -1.4966 | 0.4507 | -3.32 | **9.00e-04** | -2.0023 | 0.5706 | -3.51 | **4.51e-04** | -2.0155 | 0.5717 | -3.53 | **4.24e-04** |
| PC3 | -1.3377 | 0.5047 | -2.65 | **8.05e-03** | -0.9214 | 0.6497 | -1.42 | 1.56e-01 | -0.9332 | 0.6540 | -1.43 | 1.54e-01 |
| PC4 | 1.4997 | 0.7604 | 1.97 | **4.86e-02** | 0.9815 | 0.9600 | 1.02 | 3.07e-01 | 0.9726 | 0.9647 | 1.01 | 3.13e-01 |
| PC5 | 0.7175 | 0.9523 | 0.75 | 4.51e-01 | 0.4473 | 1.1739 | 0.38 | 7.03e-01 | 0.4317 | 1.1748 | 0.37 | 7.13e-01 |
| MC1R carrier1 | 0.0517 | 0.0191 | 2.71 | **6.81e-03** |  |  |  |  |  |  |  |  |
| MC1R carrier2+ | 0.1253 | 0.0515 | 2.43 | **1.51e-02** |  |  |  |  |  |  |  |  |
| Redhead |  |  |  |  |  |  |  |  | -0.0823 | 0.1164 | -0.71 | 4.80e-01 |
| Light blond |  |  |  |  | -0.1303 | 0.0718 | -1.81 | 6.97e-02 |  |  |  |  |
| Dark blond |  |  |  |  | -0.1170 | 0.0566 | -2.07 | **3.87e-02** |  |  |  |  |
| Light brown |  |  |  |  | -0.1334 | 0.0546 | -2.44 | **1.45e-02** |  |  |  |  |
| Dark brown |  |  |  |  | -0.1230 | 0.0540 | -2.28 | **2.29e-02** |  |  |  |  |
| Black |  |  |  |  | -0.1297 | 0.0896 | -1.45 | 1.48e-01 |  |  |  |  |
| Redhead x Sex |  |  |  |  |  |  |  |  | 0.2597 | 0.1300 | 2.00 | **4.58e-02** |

**S3 table:**
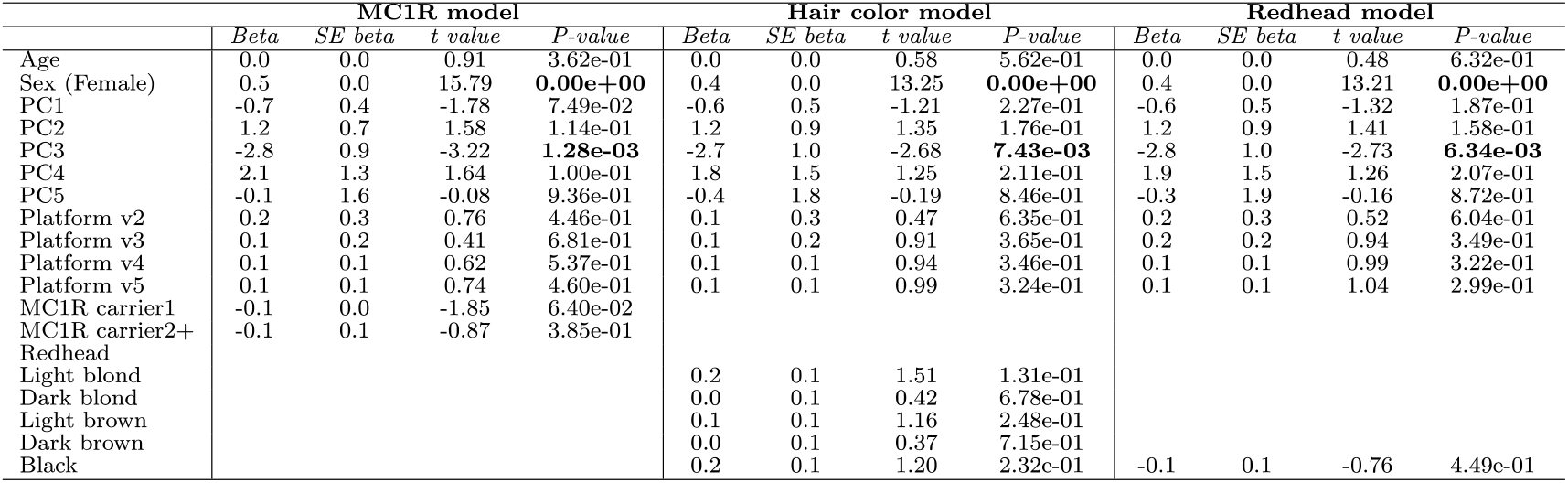
Survival models on CPT duration analyzing MC1R carriers and hair colors.

**S4 table:**
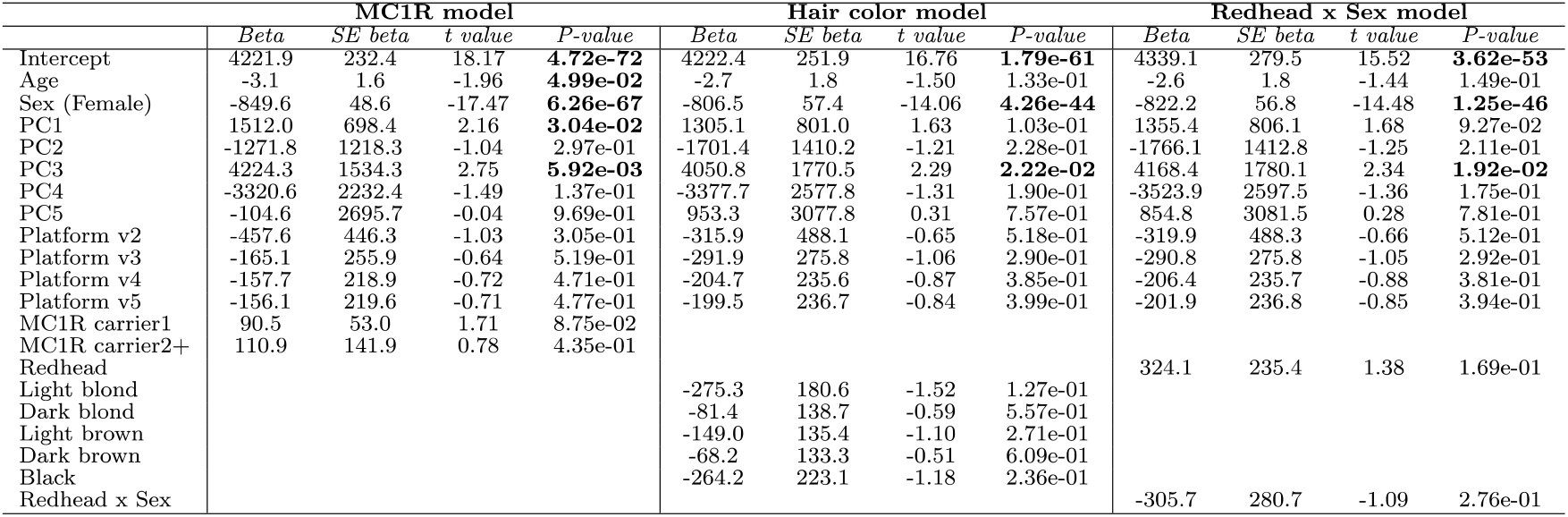
ANOVA models on CPT duration (converted in ranks), analyzing MC1R carriers and hair colors.

**S5 table:**
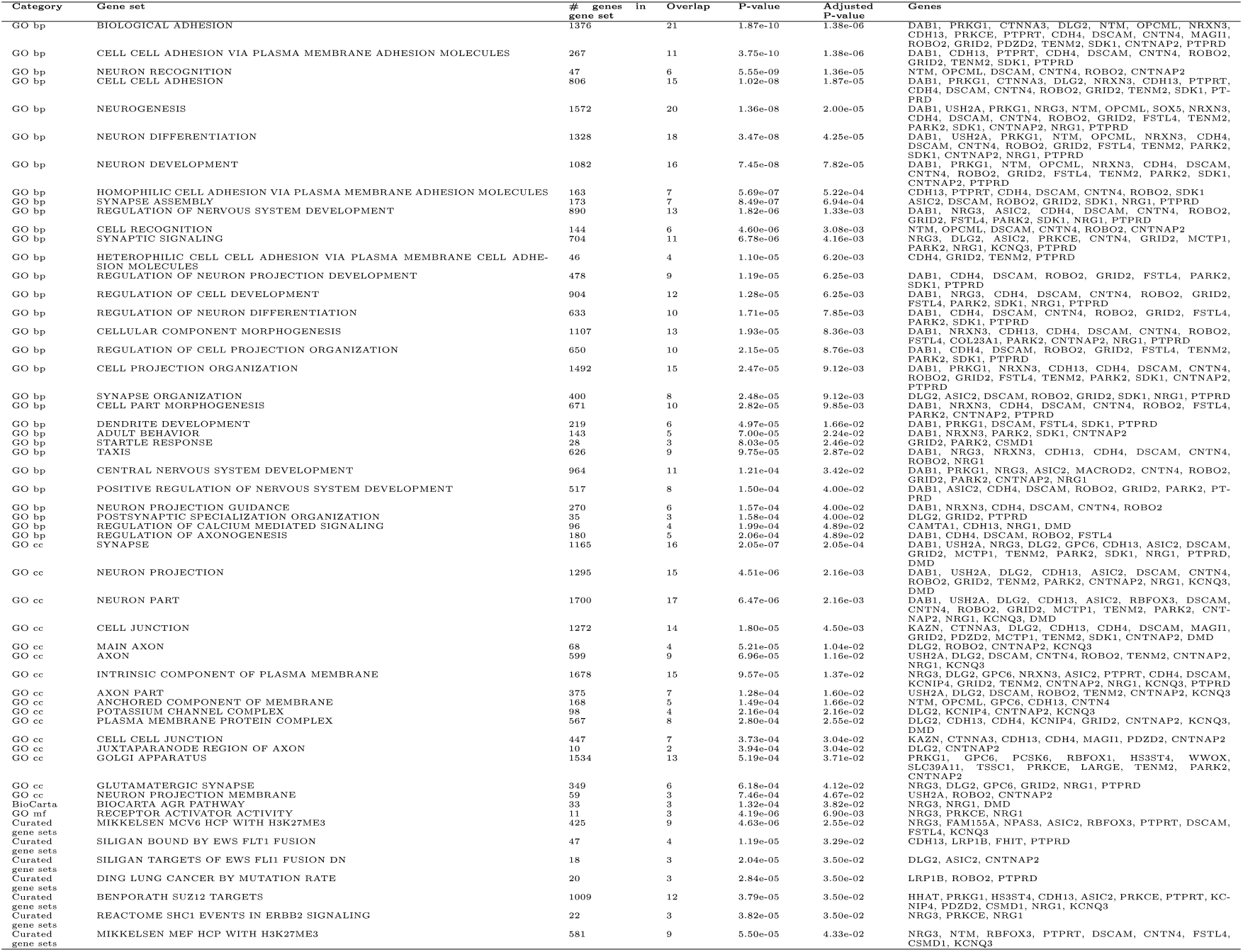
Pathway enrichment results for PSQ.

**S6 table:**
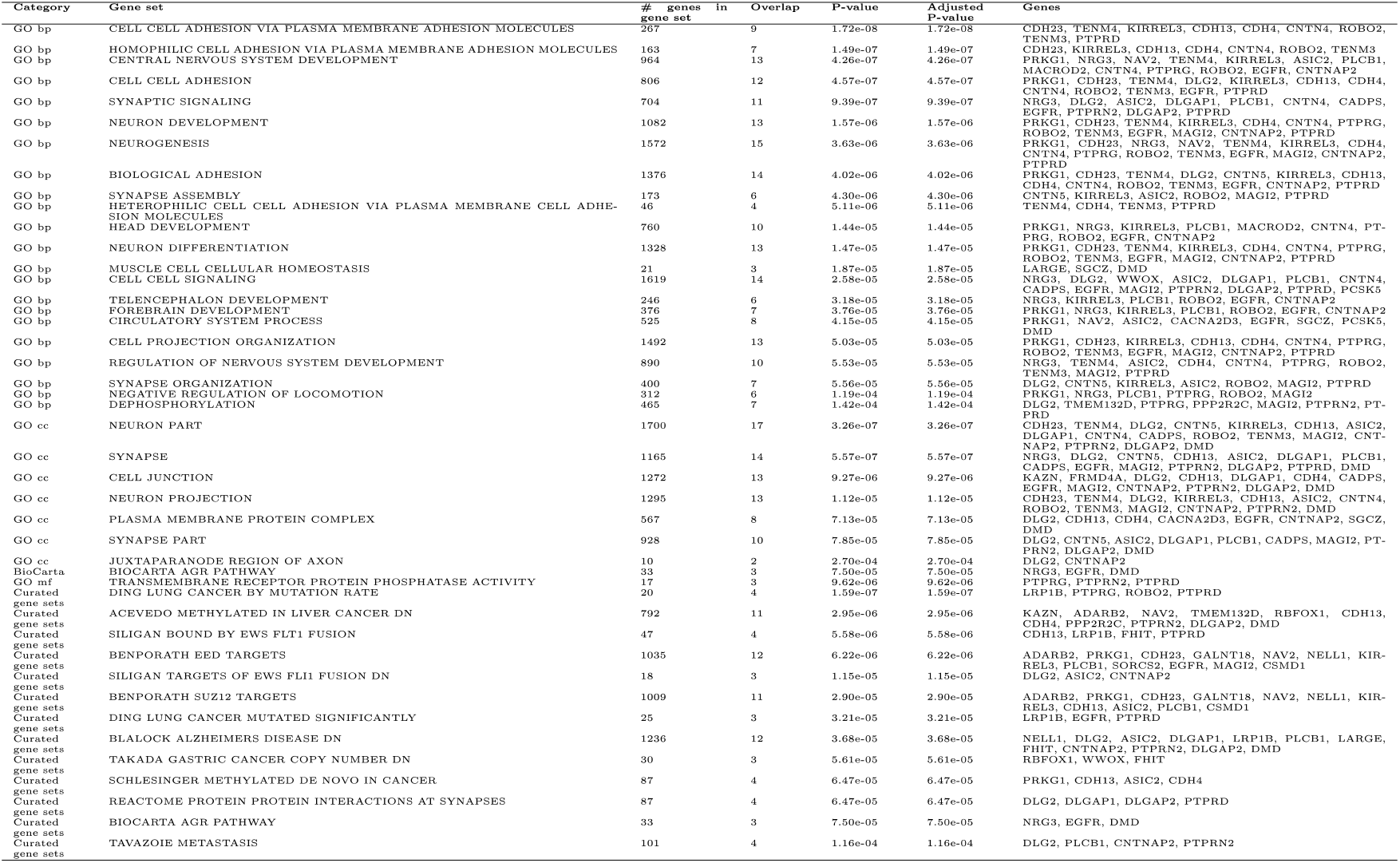
Pathway enrichment results for CPT.

**S7 table:** Gene-based (MAGMA) results for 21 nociception genes. The list of nociception genes was obtained from https://humanpaingenetics.org/hpgdb/

| Gene | Position | # of variants | PSQ |  | # of variants | CPT |  |
| --- | --- | --- | --- | --- | --- | --- | --- |
|  |  |  | P-value | Rank |  | P-value | Rank |
| ABCB1 | 7:87133179-87342639 | 22 | 4.43e-03 | 3649 | 53 | 2.55e-03 | 3101 |
| ADRB1 | 10:115803806-115806667 | 1 | 2.58e-02 | 8264 |  | NA |  |
| ADRB2 | 5:148206156-148254628 | 4 | 8.08e-03 | 4863 | 22 | 1.23e-03 | 2212 |
| ARRB2 | 17:4613789-4624795 | 4 | 1.80e-02 | 7074 |  | NA |  |
| ARVCF | 22:19954126-20004325 | 14 | 1.30e-03 | 2094 | 8 | 2.25e-02 | 8062 |
| BDNF | 11:27676440-27743605 | 4 | 1.70e-03 | 2362 | 30 | 1.77e-03 | 2585 |
| COMT | 22:19929263-19957498 | 15 | 3.44e-03 | 3232 | 3 | 1.84e-02 | 7376 |
| DRD3 | 3:113847499-113918254 | 45 | 1.13e-02 | 5714 | 31 | 9.15e-03 | 5469 |
| EDNRA | 4:148402069-148466106 | 26 | 1.53e-02 | 6566 | 4 | 3.14e-03 | 3376 |
| FAAH | 1:46859939-46879520 | 14 | 1.14e-02 | 5723 | 3 | 5.66e-02 | 11647 |
| IL1A | 2:113531492-113542971 | 1 | 8.80e-02 | 12944 | 1 | 5.97e-02 | 11851 |
| MPDZ | 9:13105700-13279660 | 33 | 5.80e-04 | 1508 | 14 | 2.72e-03 | 3192 |
| OPRD1 | 1:29138654-29190208 | 34 | 4.26e-02 | 10209 | 20 | 5.32e-02 | 11389 |
| OPRK1 | 8:54138276-54164257 | 2 | 3.87e-02 | 9835 | 26 | 1.01e-03 | 2046 |
| OPRM1 | 6:154331631-154568001 | 32 | 1.76e-03 | 2400 | 51 | 2.04e-02 | 7708 |
| SCN10A | 3:38738837-38835501 | 23 | 5.14e-03 | 3894 | 34 | 5.01e-02 | 11140 |
| SCN11A | 3:38887260-38995136 | 34 | 4.22e-02 | 10165 | 106 | 1.12e-02 | 5961 |
| SCN9A | 2:167051695-167232497 | 8 | 4.86e-03 | 3799 | 20 | 6.61e-04 | 1721 |
| TACR1 | 2:75273590-75426645 | 47 | 3.49e-03 | 3260 | 74 | 3.08e-04 | 1234 |
| TRPA1 | 8:72933486-72987819 | 19 | 6.23e-03 | 4296 | 4 | 8.66e-04 | 1913 |
| TRPV1 | 17:3468740-3512705 | 44 | 1.22e-04 | 796 | 64 | 8.46e-05 | 774 |

**S1 figure:**
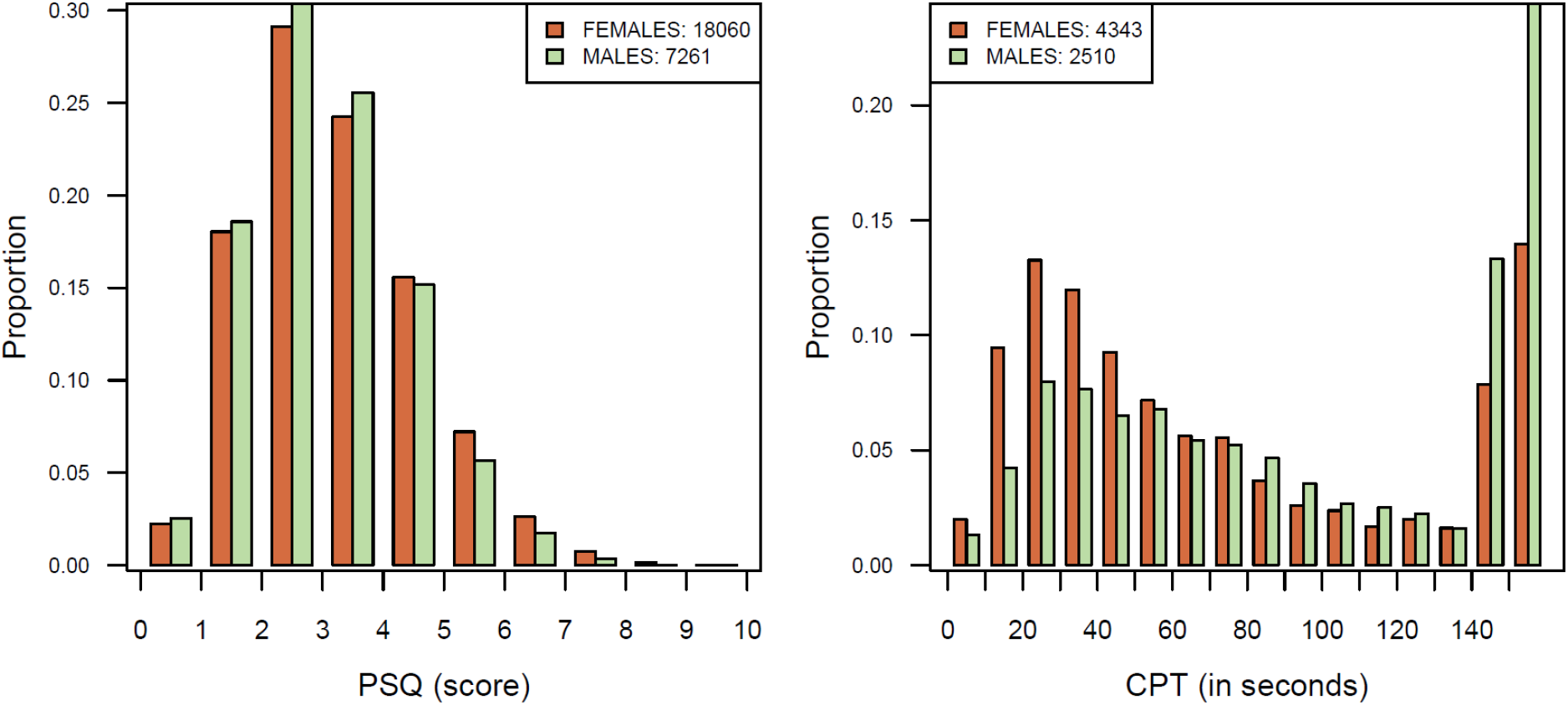
PSQ and CPT distributions.

**S2 figure:**
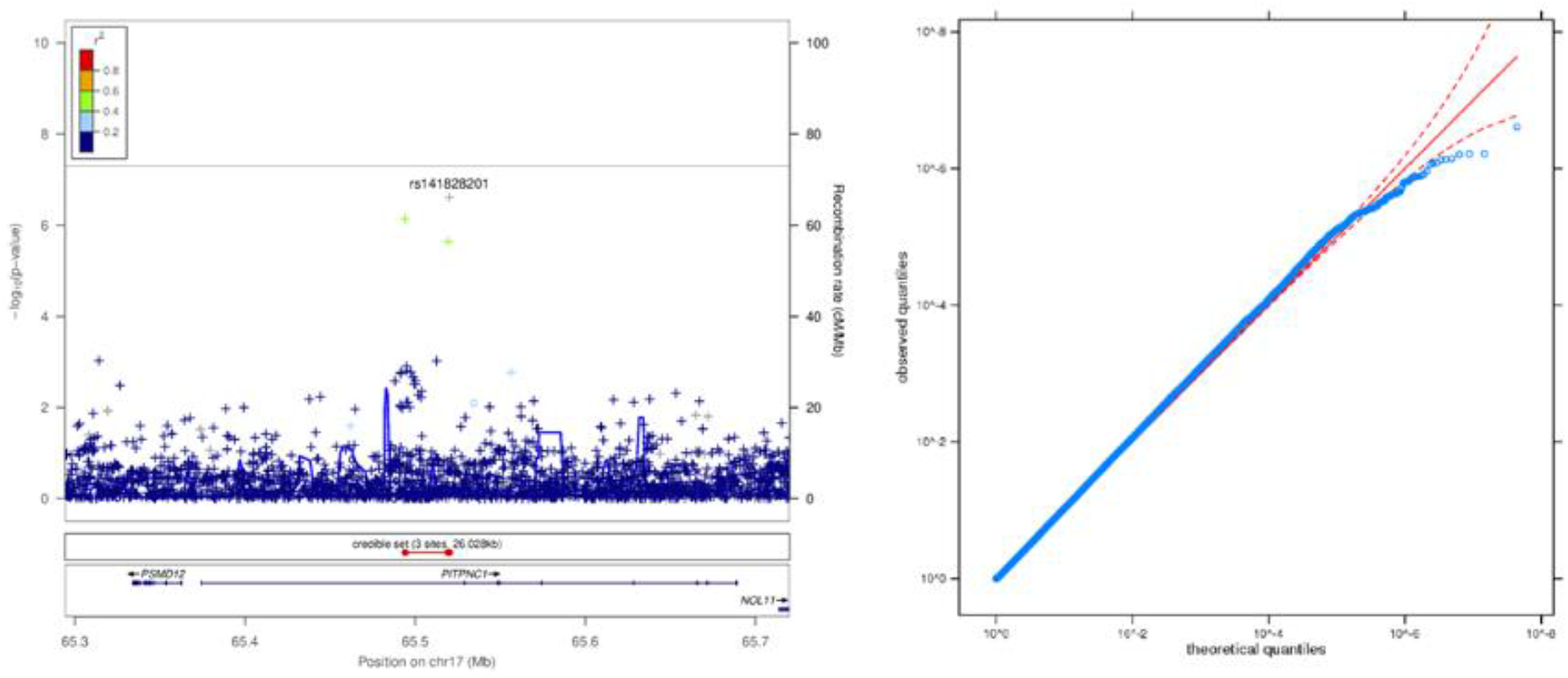
Regional association plot of the top associated loci and QQ plot from the CPT GWAS.

**S3 figure:**
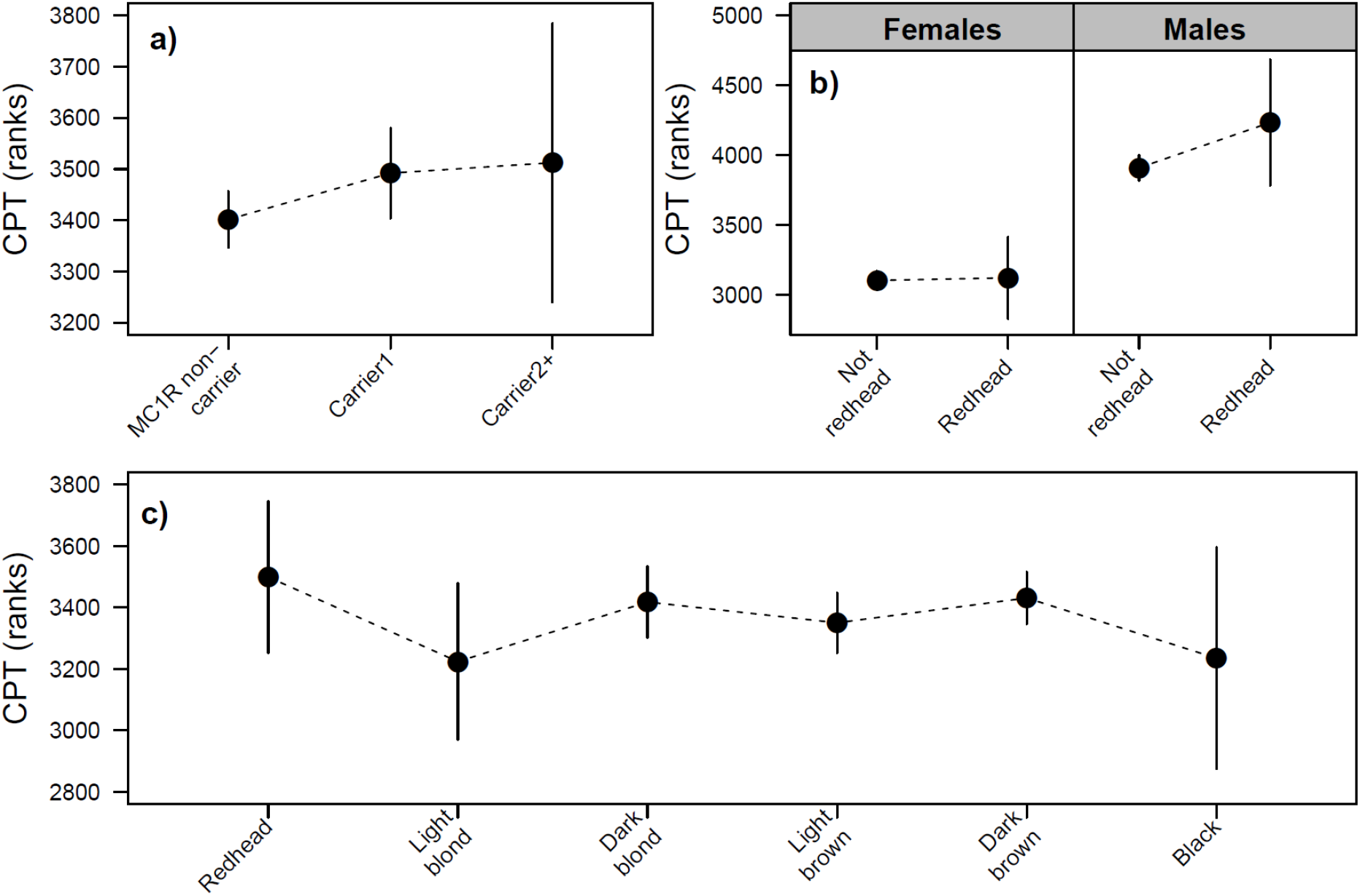
Results of MC1R and hair color models for CPT duration converted in ranks.

**S4 figure:**
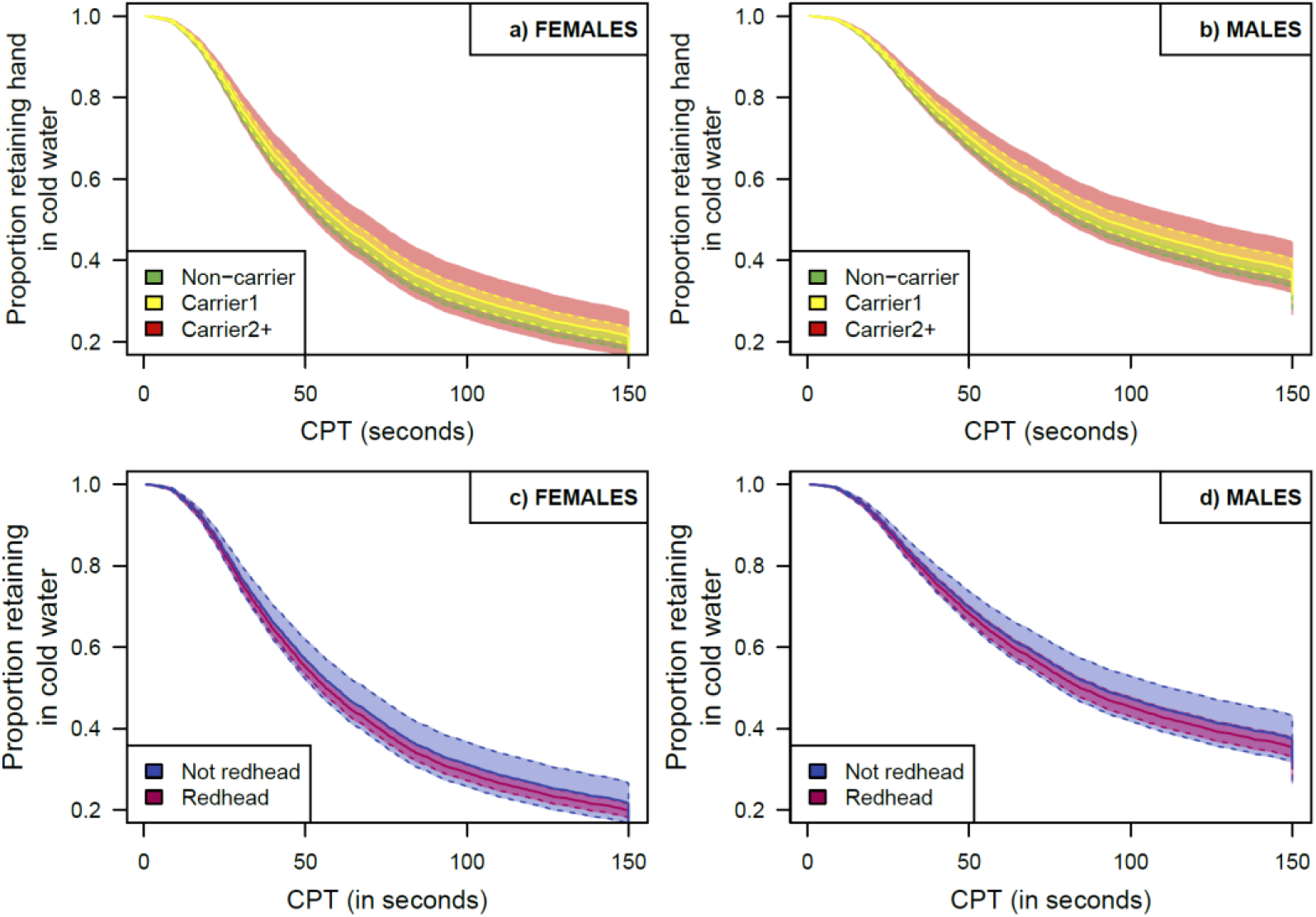
Kaplan-Meier curves for MC1R and hair color survival models on CPT duration.

**S5 figure:**
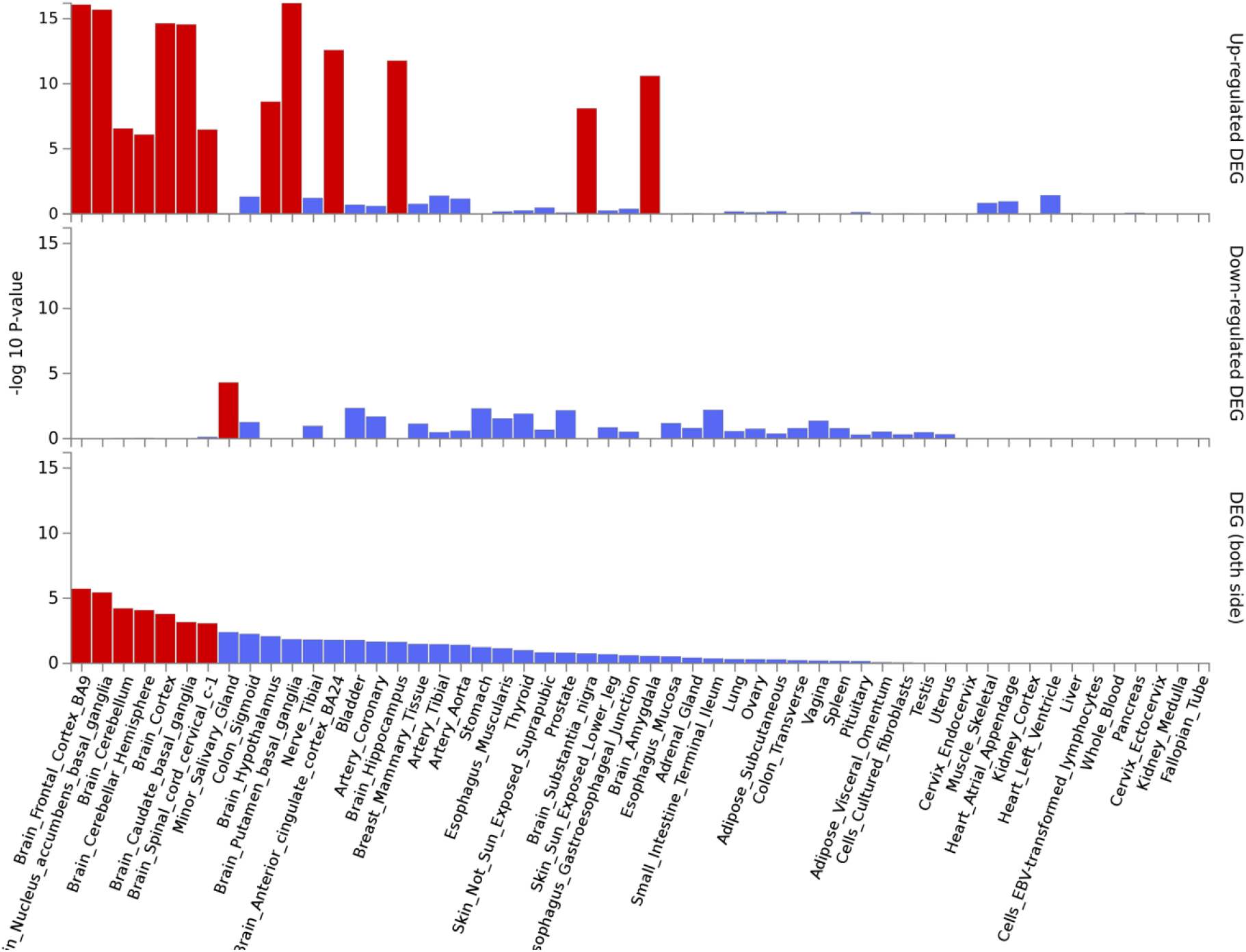
Tissue expression enrichment analysis for PSQ, based on the 58 genes in Table 3 (GTEx v8).

**S6 figure:**
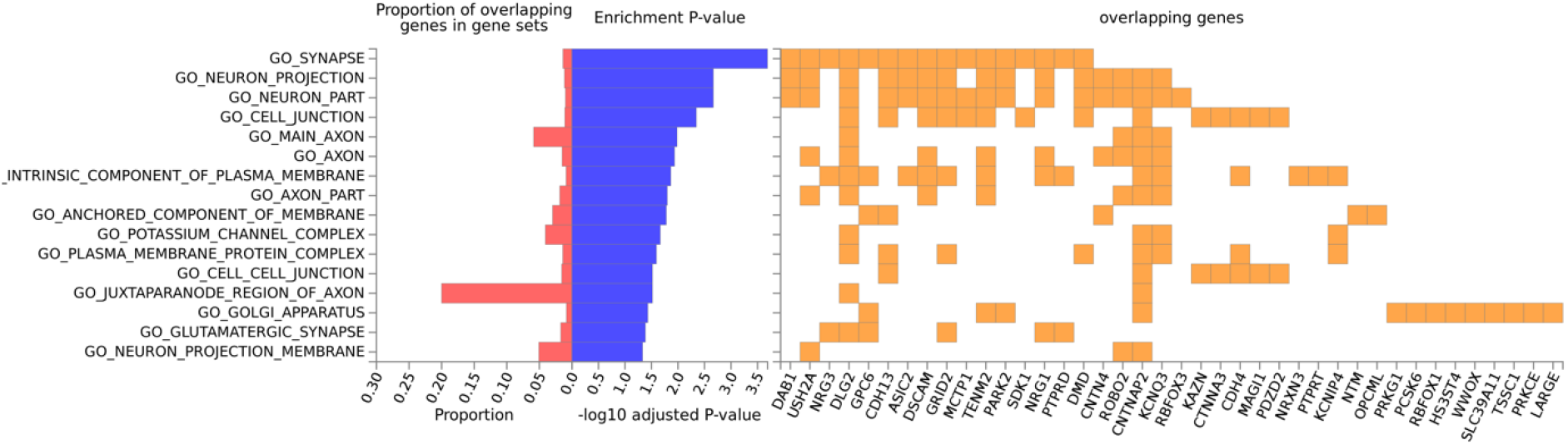
Enrichment analysis of cellular components for PSQ, based on the 58 genes in Table 3.

**S7 figure:**
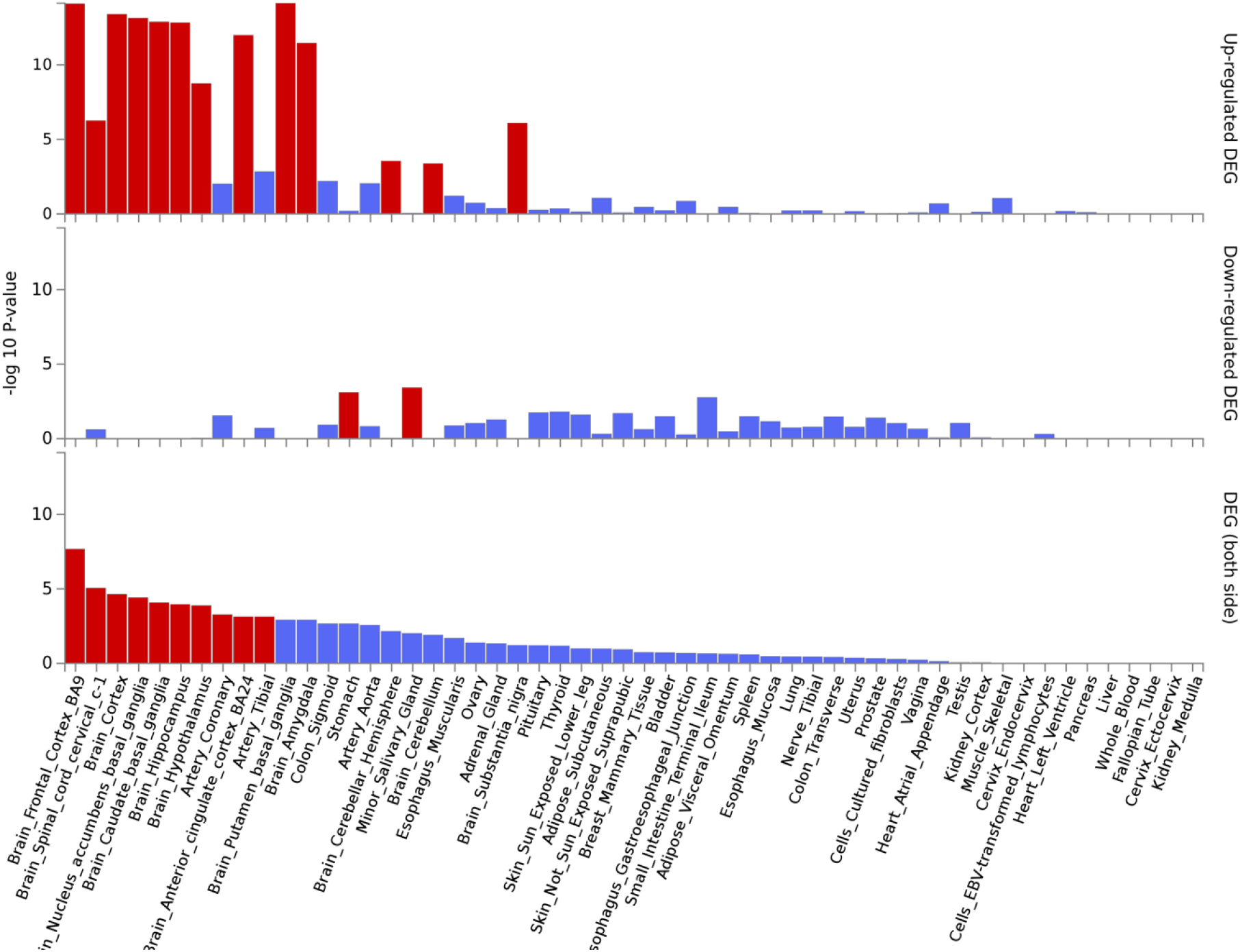
Tissue expression enrichment analysis for CPT, based on the 50 genes in Table 3 (GTEx v8).

## Notes

### Competing Interest Statement

This research was conducted by 23andMe and Grunenthal GmbH. All authors are employees of these companies.
Pierre Fontanillas and Joyce Tung are employed by and hold stock or stock options in 23andMe, Inc

